# P53 supresses transcription of the p300-E2F1-dependent gene subset by maintaining KDM5B associated with gene promoters

**DOI:** 10.1101/2025.08.25.672089

**Authors:** Karolina Gronkowska, Kinga Kołacz-Milewska, Sylwia Michlewska, Agnieszka Robaszkiewicz

## Abstract

**Background:** p53 is a transcription activator or repressor that acts mainly by having direct control over the expression of CDK inhibitor – p21 in response to DNA damage.

**Methods:** We used qPCR, Western Blot, protein co-immunoprecipitation, chromatin immunoprecipitation, RNA-seq, confocal microscopy, flow cytometry and resazurin assay to investigate how the p53 regulate gene expression upon sub-lethal doses of cisplatin.

**Results:** In this study, molecular evidence was provided for the occurrence of p53 at the subset of E2F1-driven promoters and their suppression, despite the co-occurrence of p53 with p300. P53 repressed promoters were characterized by relatively high nucleosome density and demethylation of H3K4, followed by low H3K27 acetylation and trimethylation of H3K4. Induction of the ATM/ATR-Chek1/2-p53 pathway by sub-lethal doses of cisplatin caused the release of p53 from gene promoters, chromatin relaxation and the gain of transcription permissive histone marks. Mechanistically, p53 maintained the KDM5B that is associated with gene promoters, thereby conditioning the demethylation of H3K4me3. P53 formed an immunoprecipitable complex with KDM5B, E2F1, p300 and H3K4me2 in intact cells, which decomposed with cisplatin and substantially increased the level of H3K4me3 in the p300 interactome. The extrusion of KDM5B from the chromatin was triggered by cisplatin, transient p53 silencing or KDM5B inhibition, also enabled p300 enrichment and increased gene transcription. The molecular and functional interdependence between p53 and KDM5B was observed in distinct cancer cell types, and the co-expression of *TP53* and *KDM5B* can be considered as a doxorubicin response biomarker.

**Conclusions:** p53 directly suppressed the subset of E2F1-driven genes in proliferating cells by maintaining KDM5B associated with gene promoters and inhibiting p300-mediated transcription.

## 1. Background

Tumor suppressor p53 is best known as a transcription factor that controls cell fate after multiple stress signals, including DNA damage, oncogenic stress, replication stress, oxidative stress, hypoxia and ribosomal stress. The core set of the p53 known target genes have diverse biological functions such as apoptosis, cell cycle arrest, differentiation, metabolism, autophagy and DNA repair [1]. p53 is a transcription factor that binds to the genome by its DNA recognition motif, the p53 response element (p53 RE), and facilitating or preventing the formation of active transcription complexes thereby activating or repressing genes [1,2]. Despite protracted studies on this protein, there has not been a proven universal model of p53-dependent gene activation or suppression, with these 2 opposed impacts seeming to be conditional and signal-specific. The sequence differences in p53REs were thought to determine gene induction or repression, but later genome-wide data has demonstrated that p53 is actually associated with transcriptional activation at the binding sites [2]. The repression of genes transcriptionally controlled by the E2F family of TFs is frequently assigned to indirect p53-dependent transactivation of p21 (*CDKN1A*), which inhibits CDK and stops mitotic divisions by recruiting the transcriptional repressor Rb to E2F1-driven promoters. p53 also directly transactivates *E2F7* that encodes a repressive member of the E2F family. p53 also activates the expression of various miRNAs and long non-coding RNAs, which in turn impacts transcription yield [1,3]. The analysis of the p53 direct transcriptional program using Global Run-On sequencing (GRO-seq) showed that p53 represses a subset of its activation targets (e.g. *PTP4A1*, *HES2*, *GJB5*) before its activation in the steady state of proliferating cells [4]. However, many concerns have been raised regarding the mechanism by which p53 is localized to the repressed promoters and how it switches from activation to repression. The co-repressor interaction with chromatin remains controversial, despite good experimental evidence. The tight link between the p53 mode of action and enhancer activation may explain the lack of perfect overlap between p53-dependent genes among various cell types and conditions, often referred to as proof of misinterpreted p53 targets.

Since p53 peaks are usually smaller on repressed genes when compared to activated ones, Peuget and Selivanova suggested that the low occupancy of p53, probably at the repressed promoter, reflected a more transient binding mode that was necessary to establish the repressive chromatin landscape but not to maintain it [1]. Furthermore, p53-repressed genes frequently displayed lower affinity p53 RE in their promoters, whereas most cell types which are independent of the type of activating stimuli largely comprise of high affinity p53 RE. Regarding the p53 repressive role and its interaction with DNA in a steady state, we identified and characterized some of the p53 repressed gene promoters in proliferating cells. MCF7 breast cancer cells were used as they express wild-type p53. These cells were characterized by a p53-dependent resistance to cisplatin, proved by the sensitization of the cells to drugs after the disruption of p53 by human papilloma virus (HPV) infections [5]. To avoid the complexity caused by cell cycle inhibition and likely impact of E2F-Rb at gene transcription, the cells were exposed to cisplatin at a dose which induced the arrest of mitotic divisions or cell death, but at the same time was high enough to activate ATM/ATR-Chek1/2-p53 pathway and trigger the redistribution of p53 in the genome.

The cytotoxic effect of the anticancer drug cisplatin arises through the formation of intra- and inter-strand DNA and DNA-protein crosslinks blocking the progression of DNA replication during the S-phase and thereby causing double-strand breaks (DSBs) that gave rise to insertions, deletions or even translocations [6,7]. While ataxia telangiectasia mutated (ATM) was activated by DSBs, ATM-related (ATR) was activated by stalled DNA replication forks after cisplatin treatment [7]. Phosphorylation of downstream targets, including Chek1/2 kinases and p53, lead to p21-mediated arrest in G1, whereas the phosphorylation of H2Ax at sites of DNA lesions enabled the recruitment of DNA repair proteins [8]. Although treatment conditions were chosen to avoid cell cycle inhibition, the chromatin rearrangement at the site of p53 binding in a steady state was observed. Previous reports linked p53-dependent gene suppression with the recruitment of corepressors such as histone deacetylases (HDACs), to inverted (head-to-tail) or imperfect p53REs that imparted repressive activities on p53 and enhanced competition with transcriptional activators by p53 [1,3]. Several factors were suggested that cooperated with p53 mediated repression, including NF-Y, mSin3a and HDAC1. The competition of p53 with SP1 for the repression of genes encoding telomerase and nestin was also reported [1]. In this paper we provided experimental, mechanical evidence on the role of p53 in KDM5B-mediated chromatin remodelling and repression of genes that were activated by p300 and characterized by motifs for the E2F family of transcription factors in their promoters. Attention was paid to any functional p53-p300 crosstalk, as p53-p300 interaction triggered by cisplatin has previously been described where p53 emerged as being required to recruit p300 to the promoter of *ABCC10* and hence, served as a gene activator [9]. CITED2/p300 recruitment by p53 to the promoter of the DNA repair gene *ERCC1* in response to cisplatin facilitated DNA repair is another example [7]. In this current model, the cisplatin-induced extrusion of p53 from the promoters of some E2F-dependent genes, which were characterized by the presence of p53 binding motifs, upregulated gene transcription in a way similar to p53 silencing. In the subset of p53 repressed genes, we observed previously undescribed relationships between p53 interaction with DNA, lysine acetylation and methylation of histone H3, resulting from the physical and functional interaction between p53, p300 and KDM5B.

## 2. Methods

### 2.1 Materials

Breast cancer cell line MCF7 and non-small cell lung cancer cells A549 was purchased from ATCC. DMEM High Glucose w/ L-Glutamine w/ Sodium Pyruvate, fetal bovine serum and antibiotics (penicillin and streptomycin) were from Biowest (CytoGen, Zgierz, Poland). Resazurin sodium salt (#R7017), DAPI (#10236276001) cisplatin (#232120), KU-60019 (SML1416) and sodium butyrate (#303410) were from Sigma Aldrich (Poznan, Poland). C646 (#10549), PBIT (#16272), GSK-LSD1 (#16439) from Cayman Chemical, were ordered in Biokom (Janki/Warsaw, Poland). siRNA Control (sc-37007) was purchased from Santa Cruz Biotechnology (AMX, Lodz, Poland). Nunc™ Lab-Tek™ Chamber Slide, oligonucleotides for Real-time PCR, TP53 Silencer Select siRNA (#106140), Lipofectamine RNAiMAX, OptiMem, Dynabeads™ Protein G, UltraPure™ Phenol:Chloroform:Isoamyl Alcohol (25:24:1, v/v), TRI Reagent™, High-Capacity cDNA Reverse Transcription Kit, SuperSignal™ West Pico Chemiluminescent Substrate, PageRuler™ Prestained Protein Ladder (#01154870), Pierce™ Protease Inhibitor Tablets (EDTA-free; PIC), Pierce™ Phosphatase Inhibitor Mini Tables, SlowFade™ Glass Soft-set Antifade Mountant (with DAPI), Phosphoserine/threonine/tyrosine polyclonal antibody (#XF342757), Goat anti-Rabbit IgG (H+L) Cross-Adsorbed Secondary Antibody, Alexa Fluor™ 546 (#A-11010), PowerUp™ SYBR® Green Master Mix, were from Thermofisher Scientific (Thermofisher Scientific, Warsaw, Poland). DNA Damage Antibody Sampler Kit (#9947), p300 (D2X6N) Rabbit mAb (#54062), p53 (7F5) Rabbit mAb (#2527), p53 (1C12) Mouse mAb (#2524), E2F1 Rabbit Ab (#3742), JARID1B (E2X6N) Rabbit mAB (#15327), PCNA (D3H8P) XP ® Rabbit mAB (#13110), Acetylated-Lysine Rabbit Ab (#9441), Mono-Methyl Histone H3 (K4) (D1A9) XP ® Rabbit mAb (#5326), Di-Methyl-Histone H3 (K4) (C64G9) Rabbit mAb (#9725),Tri-Methyl-Histone H3 (Lys4) (C42D8) Rabbit mAb (#9751), Acetyl-Histone H3 (Lys27) (D5E4) XP® Rabbit mAb (#8173), Histone H3 Rabbit Ab (#9715), Histone H3 (D2B12) XP® Rabbit mAb (ChIP Formulated) (#4620), anti-rabbit IgG, HRP-linked Antibody (#7074), Anti-mouse IgG, HRP-linked Antibody (#7076), Anti-rabbit IgG (H+L), F(ab’)2 Fragment (Alexa Fluor® 488 Conjugate) (#4412), Anti-mouse IgG (H+L), F(ab’)2 Fragment (PE Conjugate) (#59997), were from Cell Signaling Technologies (LabJOT, Warsaw, Poland). NEBNext® Ultra™ II DNA Library Prep with Sample Purification Beads (#E7104), NEBNext® Poly(A) mRNA Magnetic Isolation Module (#E7490), NEBNext Ultra II RNA First Strand Synthesis Module (#E7771), NEBNext Ultra II Non-Directional RNA Second Strand Module (#E6111), NEBNext® Ultra™ II DNA Library Prep with Sample Purification Beads (#E7103) and NEBNext® Multiplex Oligos for Illumina® (Index Primers Set 3) (#E7710) were from New England Biolabs (LabJOT, Warsaw, Poland). MGIEasy PCR-Free DNA Library Prep Set and DNBSEQ-G400RS High-throughput Sequencing Kit (FCL SE100) were from Perlan Technologies (Perlan Technologies Poland, Warsaw, Poland).

### 2.2 Cells and treatment

Cisplatin resistance induction and characteristics of cisplatin-resistant A549 cells was previously described by Sobczak *et al*. [9]. MCF7 breast cancer cells and cisplatin resistant A549 cells were cultured in DMEM supplemented with 10% FBS and penicillin/streptomycin (50 U/ml and 50 µg/ml, respectively) in 5% CO2. iEP300 (C646, 5 µM), iATM/ATR (KU-60019, 5 µM), iLSD1 (GSK-LSD1, 0,1µM), iKDM5B (PBIT, 5 µM) and iHDAC (Sodium butyrate, 100 µM) were added to cells 6h or 24h before analysis (depending on the tested parameters) or 2h before treatment with cisplatin. K562 leukaemia cells were cultured in RPMI supplemented with 10% FBS and penicillin/streptomycin (50 U/ml and 50 µg/ml, respectively) in 5% CO2. iEP300 (C646, 1 µM) and iKDM5B (PBIT, 1 µM) were added to cells before analysis.

### 2.3 Gene transcription with real-time PCR

For mRNA expression evaluation, total RNA was extracted from cells using TRI Reagent™. Afterwards, mRNA was reverse transcribed with the High-Capacity cDNA Reverse Transcription Kit. The expression of selected genes was measured in Bio-Rad CFX96 C1000 Touch Real-Time system, using: PowerUp™ SYBR® Green Master Mix and the manually designed primer pairs: (*ACTB* Forward: 5’ TGGCACCCAGCACAATGAA 3’, Reverse 5’ CTAAGTCATAGTCCGCCTAGAAGCA 3’, *TBP1* Forward: 5’ CACGAACCACGGCACTGATT 3’, Reverse: 5’: TTTTCTTGCTGCCAGTCTGGAC’ *DUSP12* Forward: 5’ TCCATGCTTACCACAGGGAG 3’, Reverse: 5’: AACCCAACTTGGCACTGCAT’, *FANCI* Forward: 5’ CCACCTTTGGTCTATCAGCTTC 3’, Reverse: 5’: CAACATCCAATAGCTCGTCACC’) according to the protocol provided by the manufacturer. mRNA level of particular genes was first normalized to housekeeping genes (ACTB, TBP1). The ratio between the studied and housekeeping genes was assumed to be 1 for control cells.

### 2.4 Protein detection with western blot

For protein expression evaluation cells were lysed in RIPA buffer (supplemented with 1 mM PMSF, Phosphatase Inhibitor and PIC) and sonicated (Bandelin Sonopuls HD2070); Next, proteins were separated by SDS–PAGE, transferred into a nitrocellulose membrane, and stained with primary antibodies (1:5000) at 4°C overnight. After subsequent staining with HRP-conjugated secondary antibodies (1:5000 for antirabbit and 1:2500 for anti-mouse antibodies; room temperature; 2 h), the signal was developed with the SuperSignal™ West Pico Chemiluminescent Substrate and pictures were acquired using ChemiDoc-IT2 (UVP, Meranco, Poznan, Poland). H3 was used as the control.

### 2.5 Confocal imaging

For the confocal imaging of chromatin associated proteins, cells were seeded and treated with inhibitors and cisplatin on a Nunc™ Lab-Tek™ chamber slide. Cells were fixed with a 1% formaldehyde solution in PBS at room temperature for 15 min, permeabilized and blocked with 1% FBS solution in PBS with 0.1% TritonX-100 at room temperature for 1h. Primary antibodies (1:400) were added in 1% BSA solution in PBS with 0.1% TritonX-100 and incubated at 4 °C overnight. Next, a secondary antibody (1:400) was added in 1% BSA solution in PBS with 0.1% TritonX-100 at room temperature for 2h. After washing, the slides were mounted with SlowFade™ glass soft-set antifade mountant (with DAPI). TCS SP8 (Leica Microsystems, Germany) with a 63x/1.40 objective (HC PL APO CS2, Leica Microsystems, Germany) was used for sample visualization. The samples were imaged with the following wavelength values for excitation and emission: 485 and 500-550 nm for Alexa Fluor® 488, 550 and 570-580 nm for Alexa Fluor® 546, 480 and 570-580 nm for R-phycoerythrin (PE), 405 and 430-480 nm for DAPI. The fluorescence intensity and colocalization was determined in arbitrary units (a.u.) with Leica Application Suite X (LAS X, Leica Microsystems, Germany). At least 5 fields of view were used to measure the average protein level and colocalization.

### 2.6 Co-immunoporecipitation

After treatment with inhibitors and cisplatin cells were washed three times with PBS and lysed on ice in IP buffer composed of 20 mM HEPES – KOH, 50 mM KCl, 5 mM MgCl2, 0.2 mM EDTA, 20% glycerol, 0.1% NP-40 and protease inhibitors; sonicated with the ultrasonic homogenizer Bandelin Sonopuls (HD 2070; 10 impulses, 60%); and centrifuged (3 000 rpm, 4 °C, 10 min). Supernatant was incubated with anti-p53, anti-E2F1 or anti-EP300 antibodies and control IgG at 4 °C for 2 h. For another 1 h, lysates were added with Dynabeads (10 µL); then, they were washed 5× with the IP buffer and suspended in RIPA buffer (supplemented with 1 mM PMSF and PIC) with 5% β-mercaptoethanol and gel loading buffer and heated at 70 °C for 10 min. Beads were collected on a magnetic stand and supernatant was separated by SDS-PAGE electrophoresis followed by transfer of proteins on nitrocellulose membranes. Interacting partners were detected on nitrocellulose membranes after overnight staining with corresponding primary antibodies (1:5000; 4°C; overnight) and subsequent staining with HRP-conjugated secondary antibodies (1:5000; room temperature; 2h). The signal was developed with the SuperSignal™ West Pico Chemiluminescent Substrate and pictures were acquired using ChemiDoc-IT2 (UVP, Meranco, Poznan, Poland).

### 2.7 Chromatin immunoporecipitation (ChIP)

Chromatin immunoprecipitation was carried out according to the protocol previously described [10]. DNA was extracted using phenol:chloroform:isoamyl alcohol (25:24:1). P53 was immunoprecipitated with p53 (7F5) Rabbit mAb, EP300 with p300 (D2X6N) Rabbit mAb, KDM5B with JARID1B (E2X6N) Rabbit mAB, H3K4me3 with Tri-Methyl-Histone H3 (Lys4) (C42D8) Rabbit mAb and H3K27ac with Acetyl-Histone H3 (Lys27) (D5E4) XP® Rabbit mAb in non-treated MCF7 cells and treated with cisplatin/inhibitors/siRNA MCF7 cells.

1μg of immunoprecipitated DNA fragments was converted into library for sequencing using NEBNext® Ultra™ DNA Library Prep Kit with Sample Purification Beads for Illumina® and NEBNext® Multiplex Oligos for Illumina® (Index Primers Set 3) according to instruction provided by manufacturer. DNA library was sequenced on NextSeq 550 in the Department of Clinical Genetics, Medical University of Lodz, and data were released as fastq files.

Histone modifications and transcriptional regulators on DUSP12 and FANCI promoters was analyzed using ChIP-qPCR using PowerUp™ SYBR® Green Master Mix, 0.1% DMSO and manually designed primer pairs. Primers were designed based on p53 binding motifs from ChIP-seq data : *FANCI* Forward 5’ CAGGAGGGAAGCTGAACCTG 3’, Reverse 5’ ATGAAGACTGAAGGGGTGCC 3’; *DUSP12* Forward: 5’ CCAAGAGAGGAGGAGGGTTG 3’, Reverse: 5’ CAATAATCGGGGTGGGTGTCT 3’, Data were normalized to samples containing non-specific IgG or histone H3 (for H3K4me3 and H3K27ac).

### 2.8 Library preparation for RNA-seq

Total RNA was isolated using TRI Reagent™. Next, 10 ng of DNA-free total RNA was used to isolate mRNA using NEBNext® Poly(A) mRNA Magnetic Isolation Module.Next mRNA was converted do cDNA using NEBNext Ultra II RNA First Strand Synthesis Module and NEBNext Ultra II Non-Directional RNA Second Strand Module according to manufacturer instructions. The purified cDNA was used for RNAseq library preparation using MGIEasy PCR-Free DNA Library Prep Set according to manufacturer’s protocol. Sequencing was performed using DNBSEQ-G400 in the Department of Clinical Genetics, Medical University of Lodz, and data were released as fastq files.

### 2.9 Bioinformatic analysis in usegalaxy.org (server version 25.0.2.dev0)

After the quality control in FastQC the raw reads (as fastq.gz) derived from ChIP- and RNA-Seq were trimmed to reads with Q_phred_>30.

ChIP-Seq-derived reads were mapped to human genome (hg19) with Bowtie2. For peak calling we use MACS2 callpeak with peak detection based on p<0.001. The genome coverage was normalized to counts per million (CPM) with bamCoverage and saved in bigwig for further visualization in UCSC Human Genome Browser. Gene promoters were assumed as regions ± 5kbp from transcription start sites (TSS), which were derived from UCSC GTEx Gene V8 (Table: gtexGeneV8). Heatmap and plot distribution of proteins were generated by computeMatrix with bigwig as score files, regions to plot in bed from MACS2 peaks or USCS tables, a reference-point (center of region) and selected distance upstream and downstream. Pearson correlation coefficient and correlation heatmap for E2F1 motifs in the genome (filtered out from UCSC table wgEncodeRegTfbsClusteredV3) and gene promoters (from Table: gtexGeneV8). Overlapping region in genome were searched with Intersect tool for data saved in intervals or bedtools Intersect intervals for data saved in bed. The coverage with mapped reads (as BAM) for specific genomic regions or promoter subsets (as BED) and considered features (proteins or H3 modifications) was counted with bedtools MultiCovBed. The FASTA sequences for p53 peaks in activated and repressed promoters (as BED) were derived from server indexed file (fasta id: Human (Homo sapiens): hg19) using bedtools getfasta. P53 binding motif was taken from JASPAR (MA0106.3) and aligned to FASTA sequences in MEME Suite 5.5.8 tool: Find Individual Motif Occurences (FIMO). Top20 transcription factors and co-factors for the promoter subsets and, then, for p300-p53 promoters were identified and counted by comparing promoter regions with wgEncodeRegTfbsClusteredV3 table (as BED), and transcription factors were sorted by frequency of their occurrence.

RNA-Seq-derived data were mapped to hg19 reference genome with HISAT2 with default parameters, and gene expression was measured by featureCounts using built-in hg19 genome. Differential gene expression analysis and normalization (filtered and normalized counts tables) were carried out with limma-voom using the following parameters: normalization method – TMM; filtering on count-per-million (CPM) with CPM below 1 in minimum 3 samples; minimum Log2 fold change – 1; p value adjusted threshold – 0.05; p value adjustment method – Benjamini and Hochberg (1995). Normalized counts were then plotted using Heatmap2 and the following parameters: compute z score on columns; clustering rows and not columns; distance method – Euclidean; clustering method – average (UPGMA).

Datasets from SRA for MCF7 cell line:

|  |  |
| --- | --- |
| H3K27ac | SRR12349339 and SRR10887027 |
| H3K4me3 | SRR12349279 and SRR10886967 |
| H3K4me2 | SRR7686906 |
| H3K4me1 | SRR7952790, SRR7952791, SRR7952792 and SRR7952793 |
| H3K9me3 | SRR14678506 and SRR14678507 |
| H3K27me3 | SRR3159919 and SRR3159920 |
| ER | SRR23071683 and SRR23071681 |
| POLR2A | SRR10320030 |
| POLR2A Ser2P | SRR10320034 |
| LSD1 | SRR14213437 |
| EZH2 | SRR15724297 |
| KDM5B | SRR827515 |
| HDAC1 | SRR14777638 |
| MLL1 | SRR4011725 |
| MLL2 | SRR4011726 |
| MLL3 | SRR3561357 |
| MLL4 | SRR19782007 |
| H3K4m3e in WT | SRR7686907 |
| H3K4m3e in KDM5B KO | SRR7686919 |

Datasets from SRA for A549 cell line:

|  |  |
| --- | --- |
| H3K27ac | SRR20067327 and SRR20067326 |
| H3K4me3 | SRR24225816 and SRR24225817 |
| H3K4me2 | SRR24225823 and SRR24225822 |
| p53 | SRR28401139 and SRR28401138 |

Datasets from SRA for A2780 cell line:

|  |  |
| --- | --- |
| H3K27ac | SRR14295681 and SRR14295682 |
| H3K4me3 | SRR8887290 and SRR8887291 |
| H3K4me2 | SRR8887294 and SRR8887295 |
| p53 | SRR8887302 and SRR8887303 |

Datasets from SRA for K562 cell line:

|  |  |
| --- | --- |
| H3K27ac | SRR2085858 and SRR2085876 |
| H3K4me3 | SRR26908600 and SRR26908586 |
| H3K4me2 | SRR26908586 and SRR26908601 |
| p53 | SRR9104908, SRR9104909, SRR9104910, SRR9104884, SRR9104885 and SRR9104886 |
| KDM5B | SRR568451 and SRR568452 |

### 2.10 Cell cycle analysis

Cells treated with different cisplatin concentrations was trypsinized and washed with PBS. Next cells were fixed with 0,5% formaldehyde solution in PBS,15 min at room temperature. In the next step cells was incubated with 0,1% Triton X-100 solution. After subsequent washing cells were treated with RNAse A solution in 1% BSA solution and incubated 2h in 37°C. After incubation cell suspension was staining with DAPI (1 ug/mL).The fluorescence intensity was measured by a flow cytometer LSR® II (Becton Dickinson) at ex: 405nm / em: 480 nm/nm. The cell population was discriminated based on FSC-A and SSC-A parameters in Flowing Software 2.

### 2.11 Resasurin toxicity assay

The day prior to treatment, cells were seeded at a density of 2000 cells per well on Nunc® MicroWell™ 384-well optical bottom plates. After incubation with cisplatin cells were incubated with the resazurin solution (5 µM) in the growth medium at 37 °C for 2 h. The fluorescence that corresponds to the metabolic activity of living cells was measured with a fluorescence microplate reader (BioTek Synergy HTX, Biokom, Poland) at excitation 530 and emission 590 nm. The fluorescence value for control cells was assumed to be 100%.

### 2.12 Human xenografts and tumor sampling

100 000 cisplatin-resistant A549 cells suspended in Geltrex™ LDEV-Free Reduced Growth Factor Basement Membrane Matrix were injected subcutaneously into 6-week-old female athymic, immunodeficient mice (Crl:NU(NCr)-Foxn1nu; Animalab, Poland) and treated with 10 mg/kg cisplatin or PBS once a week for 6 weeks. After the treatment tumors were isolated, transferred and stored in PBS with 10% paraformaldehyde. The entire experiment with animals was performed in the Animal Facility of Faculty of Biochemistry, Biophysics and Biotechnology, Jagiellonian University, Poland, under permission nr 72/2023 issued on 06.04.2023 by the 2nd Local Ethical Committee in Krakow.

Tumors were paraffin embedded and sectioned using microtome (Leica RM2255, Leica Biosystems Inc.) to 10 µm thick slides, washed with xylene and ethanol to remove the paraffin and the Proteolytic-Induced Epitope Retrieval (PIER) method was used. Sections in trypsin solution (0.05%) were incubated 20 minutes at 37°C in humidified chamber. Next, samples were permeabilized and blocked with 1% FBS solution in PBS with 0.4% TritonX-100 at room temperature for 1h. Primary antibodies (1:400) were added in 1% FBS solution in PBS with 0.1% TritonX-100 and incubated at 4°C overnight. Next, a secondary antibody (1:400) was added in 1% FBS solution in PBS with 0.1% TritonX-100 at room temperature for 2h. After washing, the slides were mounted with SlowFade™ glass soft-set antifade mountant (with DAPI). TCS SP8 (Leica Microsystems, Germany) with a 63x/1.40 objective (HC PL APO CS2, Leica Microsystems, Germany) was used for sample visualization. The samples were imaged with the following wavelength values for excitation and emission: 485 and 500-550 nm for Alexa Fluor® 488, 550 and 570-580 nm for Alexa Fluor® 546.

### 2.13 Cell lines and clinical data analysis

Validation of the predictive co-expression of *KDM5B* and *TP53* to response of 581 cancer cell lines to doxorubicin was prepared in Rocplot.org [11]. The same method was applied to generate interdependence between mRNA level of *KDM5B* and response of ER+ breast cancer patients to anthracycline-based chemotherapy or ovarian cancer patients to platin-based chemotherapy [12]. Dot-plot correlation graph with mRNA level of *TP53* and *KDM5B* was generated using 32 datasets TCGA PanCancer Atlas studies available in cBioPortal [13]. Overall survival of patients, which varied in *KDM5B* expression, was generated in kmplot.com [14].

### 2.14 Statistical analysis

Data are shown as mean ± standard deviation (SD). Parametric or non-parametric test was conducted after testing Gaussian distribution of data with the Shapiro–Wilk test. Student’s t test or the Mann– Whitney test was used to calculate statistically significant differences between two samples, while one-way analysis of variance (ANOVA) or Kruskal-Wallis test followed by corresponding post hoc test was carried out to compare multiple samples. Statistics were calculated using GraphPad Prism 8.01 software. Statistically significant differences were marked with * when p < 0.05, ** when p < 0.01, *** when p < 0.001.

## 3. Results

### 3.1 Activation of ATM/ATR-Chek1/2-p53 pathway by cisplatin causes redistribution of p53 in the genome

In the search for genes repressed by p53 in MCF7 cells, a range of cisplatin concentrations (0.5-80 µM) were first tested to find the correct dose of the drug which activated the ATM/ATR-Chek1/2-p53 pathway without any considerable impact on cell cycle progression. There was no cell cycle arrest for up to cisplatin concentrations of 5 µM, as shown by the expression of mitotic markers, cell cycle analysis and viability test (Fig.1A, Supplem. Fig. 1A -B). At a concentration of 5 µM the considered drug induced mild genotoxic stress was confirmed by increased DNA double strand breaks (pH2Ax) and the phosphorylation of ATM/ATR-Chek1/2-p53 cascade components, while the level of p53 remained unchanged (Fig.1A, Supplem. Fig 1C). Surprisingly, the fluorescence of p53 in the nucleus substantially declined after cisplatin, quantified by confocal imaging (Fig. 1B). This suggested that p53 is bound to DNA in a steady state and that activation of ATM/ATR-Chek1/2-p53 triggered redistribution of p53 from the chromatin, which was followed by the extrusion of a considerable portion of p53. Pharmacological inhibition of ATM/ATR prevented the p53 eviction and maintained any p53 that was bound to DNA (Fig. 1C). This indicated that p53 activation induced a decrease in p53 interaction with DNA. This was also observed in the ChIP-Seq experiment, where the total number of p53 peaks in the genome and at the gene promoters declined after cisplatin (Fig. 1D). P53 enriched regions were shifted in the genome and detected upstream closer to TSS and downstream further from TSS (Fig. 1E). p53 enriched regions were created one peak right before TSS in untreated cells, whereas after cisplatin, p53 split into 2 peaks adjacent to TSS, which may point to the formation of transcription initiation complex in the proximity to the first exon (Fig. 1F). This observation suggested that p53 is associated with repressed promoters in proliferating, untreated cells, whereas its activation with the drug moves the protein transcriptionally permissive promoters. As a control, the p53 occurrence at the promoter of CDKN1A was monitored, as a known direct target of p53 (Fig. 1G). In the steady state, p53 was found at 3 regions upstream of *CDKN1A*, with the peak localized directly at TSS considerably erased and split after cisplatin (cropped, red box). This corresponded to the p53 redistribution around TSS shown in Fig. 1F. In a Venn diagram the peak enriched regions between untreated and cisplatin treated cells in the genome were compared, specifically at the gene promoters assumed to be TSS ± 5 kbp (Fig. 1H-I). Only 26% of p53 peaks detected in untreated cells remained unchanged after cisplatin, which caused both p53 extrusion and *de novo* recruitment to chromatin. The prevailing number of mobile peaks were detected in the intergenic regions.

**Fig. 1.**
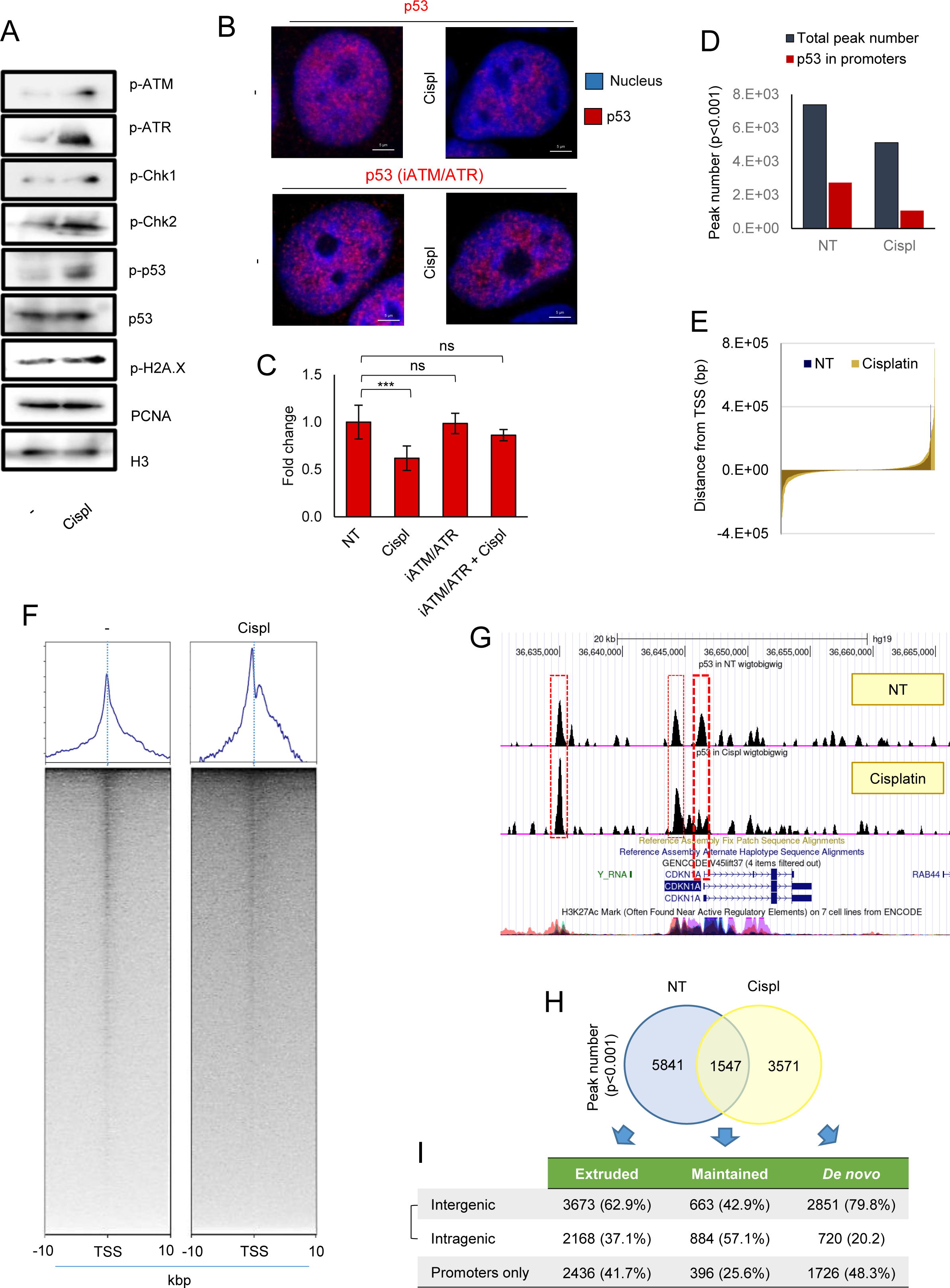
Activation of ATM/ATR-Chk1/2-p53 pathway by cisplatin induces substantial p53 extrusion from the chromatin. (A) Cisplatin causes DNA lesions and activates DNA damage response cascade as evidenced by western blot detection of phosphorylated p-H2AX and components of ATM/ATR-Chk1/2-p53 pathway. Histone H3 served as loading control. Treatment condition: 5 µM cisplatin for 6 h. (B-C) Association of p53 with chromatin declines after cisplatin but (D-E) it is prevented by iATM/ATR (KU-60019, 5 µM added 2 h prior to cisplatin). Immunostaining of p53 bound to chromatin was visualized by confocal microscopy (B), and (C) fluorescence intensity was quantified in arbitrary units (a.u.) with Leica Application Suite X. (D) Extrusion of p53 from the chromatin in response to cisplatin is confirmed by decreased p53 peak number. p53-enriched regions were scored after MACS2 peak calling at p<0.001. Gene promoters were assumed as regions ± 2 kbp from TSS. (E-F) Cisplatin triggers p53 redistribution in the genome. (E) The most significant shift in localization of p53 peaks is observed for regions furthest from TSS. (F) Cisplatin causes also considerable change in the profile of p53 occurrence around TSS. (G) In untreated cells p53 is bound to chromatin at the regulatory regions of *CDKN1A*, which are marked by massive H3K27 acetylation. The level of p53 at TSS declines after cisplatin. p53 ChIP-Seq data as bigwig are visualized in UCSC Human Genome Browser (hg19) and align to H3K27ac from ENCODE. (H) Genome-wide response of chromatin to cisplatin is followed by substantial p53 extrusion and considerably smaller number of *de novo* enriched regions. Localization of p53 peaks (p<0.001) is compared on venn diagram between untreated and cisplatin treated cells. (I) Division of peaks into specific locations in the genome (intragenic vs intergenic) and gene promoters show the strongest p53 redistribution in the intergenic regions.

These results suggest that p53 is bound to the subset of genomic regions in proliferating, untreated cells and is mostly redistributed to other locations after cisplatin.

### 3.2 p53 extrusion from p53-p300-E2F1 assembled complexes in steady state is followed by p300 enrichment at the gene promoters

In the search for possible interacting transcription co-factors , the Top 20 motifs were identified that overlap with the 3 p53 peak types: extruded, remaining and *de novo* (Fig. 2A). All 3 peak types shared 18 transcription factor motifs. p53 extruded peaks were frequently found at E2F4 and EGR1 binding sites, whereas unchanged and *de novo* deposited peaks were at p300 and RCOR1 motifs. When supported by previous studies, this observation may suggest that p53 recruitment to chromatin in response to cisplatin could activate transcription by interacting with p300. However, when p53 peaks at the gene promoters featured by p300 motifs were counted, we surprisingly found their prevailing number in untreated cells (Fig. 2B). In untreated cells, p53 peaks on p300 motifs were also featured by more frequent occurrences of the binding sites for E2F family members, specifically E2F1, E2F4 and E2F7 (Fig. 2C), as well as for chromatin remodelling enzymes such as KDM5B and HDAC1. Knowing that MCF7 cells are heterozygous for *EP300*, where one allele may harbour R1356* mutation, we first confirmed that p300 is considerably present in these cells (Supplem.Fig 1D). The confocal imaging of p53 and p300 in cell nuclei confirmed the co-localization on the chromatin of untreated cells, and declined in co-distribution after cisplatin (Fig. 2D-E). Although the total level of p53 considerably decreased in cells challenged with the anticancer drug, the p300 occurrence on chromatin substantially increased (Fig. 2F). This finding was also confirmed by a ChIP-Seq experiment for p300, where the enzyme was enriched at considered E2F1 motifs after 6 h cell treatment with cisplatin, but was erased strikingly after longer incubation (Fig. 2G). This was confirmed by confocal microscopy (Supplem. Fig 1E-G) and protein co-immunoprecipitation (Supplem.Fig 1H-I) and showed an increase in the interaction between p300 and E2F1 after treatment of the cells with cisplatin. However, as suggested by the p53 distribution on the p300 motifs (in Fig. 2B) and p53-p300 colocalization in the genome, cisplatin caused a substantial decline in the number of p53-p300 overlapping peaks at the E2F1 motifs (Fig. 2H). This indicated that p53 could interact with p300 at the promoters with E2F motifs in intact cells and that p53 *de novo* recruitment to chromatin in response to cisplatin may not be linked with co-recruitment of p300. The Pearson correlation coefficient between p53 and p300 at the gene promoters (genome wide) were similar, regardless of the cisplatin treatment (Fig. 2I), but dropped substantially at the promoters with E2F motifs from 0.6 to 0.48. Co-immunoprecipitation of p53 indicated the assembly of p53, p300 and E2F1, but cisplatin caused the decomposition of the complex and a confirmed decline in p53-p300 co-occurrence at the gene promoters with E2F1 motifs (Fig. 2J). Inhibition of ATM/ATR enhanced p53-p300-E2F1 interaction and prevented the disassembly of the complex in response to cisplatin. A prevailing number of pre-existing p53-p300 complexes enriched at E2F1 promoters were lost after cisplatin and lower numbers occurred *de novo* (Fig. 2K). Despite the extrusion of p53 from p53-p300-E2F1 promoters, p300 remained bound to the DNA after exposure of the cell to cisplatin. The quantification of p300 at the p53 extruded and *de novo* enriched promoters indicated an increase in p300 after 6 h, which was considerably higher at p53 extruded than *de novo* recruited p53 (Fig. 2L). Therefore, p53 extrusion from pre-existed p53-p300-E2F1 complexes is associated with a substantial enrichment of p300.

**Fig. 2.**
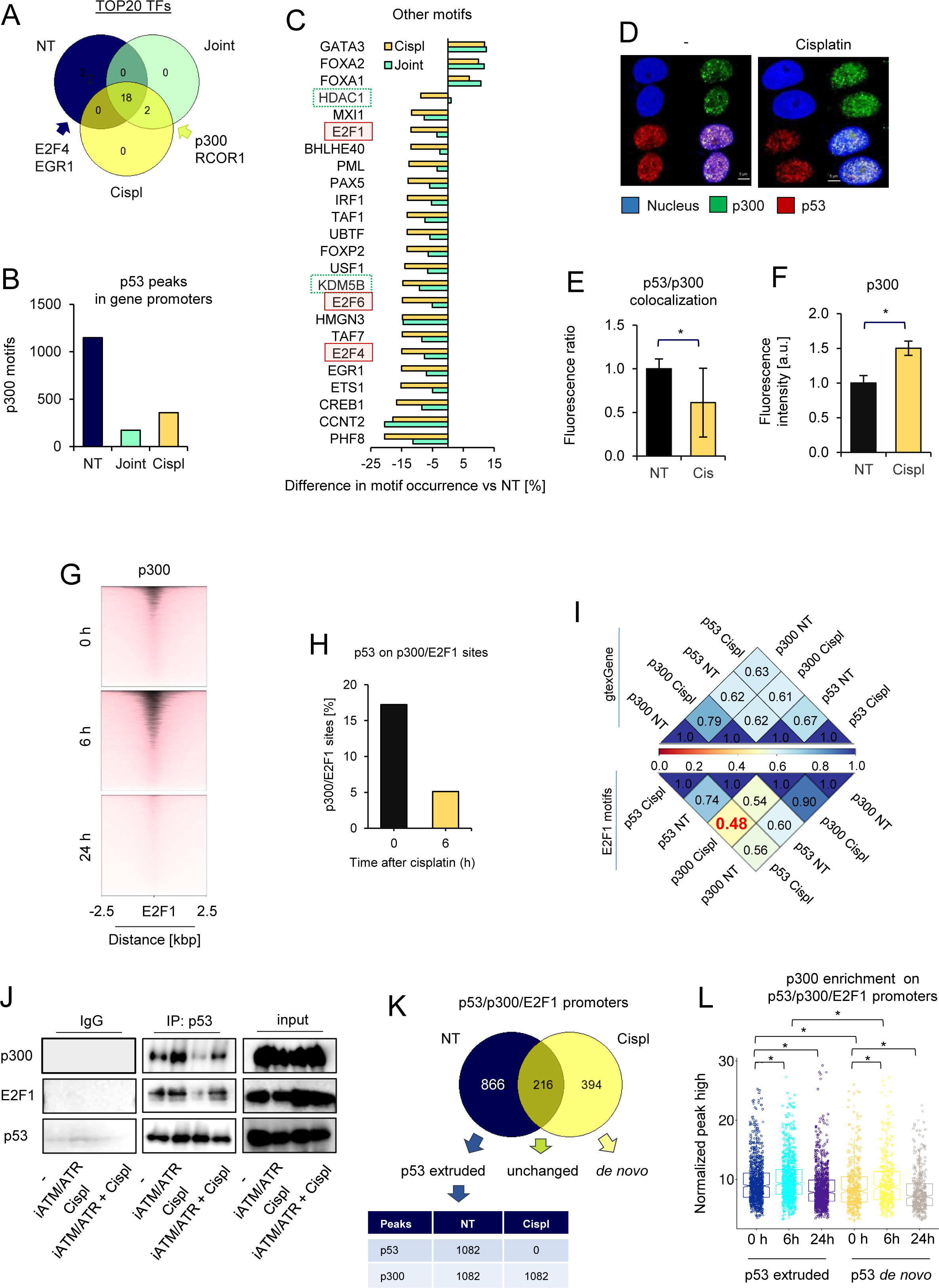
p53 extrusion from the subset of p300-bound promoters leads to significant p300 enrichment in response to cisplatin. (A)Extruded, maintained and de novo enriched p53 genomic regions were searched for transcription factors in UCSC table wgEncodeRegTfbsClusteredV3. TOP20 most frequently occurred TFs are compared on venn diagram. (B) p53 enriched region in untreated cells frequently overlap p300 binding motifs. (C) Analysis of TF binding sites indicates that p300 motifs more often co-occur with E2Fs and histone remodeling enzymes such as HDAC1 and KDM5B at p53 peaks, which are erased after cisplatin. The difference in frequency of motif occurrence at p300 motifs was compared between p53 peaks, which are extruded, maintained or deposited de novo after cisplatin. (D-E) Nuclear colocalization of p300 and p53 declines after cell treatment with cisplatin. (D) Confocal images show nuclear distribution of p300 (green – AlexaFluor488) and p53 (red – PE). DNA (blue) is stained with DAPI. (E) Quantification of fluorescence of p300 bound to DNA indicates its recruitment to chromatin in response to cisplatin. (F) Decline of red to green fluorescence ratio after cisplatin suggests spacial separation of p53 and p300. (G) p300 binds to chromatin at E2F1 motifs 6 h after cisplatin, but is substantially extruded after another 18 h. Heatmap of p300 distribution was plotted from matrix of E2F1 motifs (filtered out from UCSC table wgEncodeRegTfbsClusteredV3 table ± 2.5 kbp) as bed and p300 scores as bigwig after p300 peak calling in MACS2. ChIP-Seq for p300 was carried out after 0, 6 and 24 h cell treatment with cisplatin. (H) p53 is extruded from p300 peaks at E2F1 motifs 6 h after cell treatment with cisplatin. (I) Correlation plot shows the decline in p53 and p300 co-distribution at E2F1 motifs after cisplatin. For multiBigwigSummary, E2F1 motifs and gene promoters (± 2 kbp around TSS of gtexGene table) served as genomic regions, whereas p53 and p300 bigwig files were used for scoring. (J) p53 forms immunoprecipitable complexes with p300 and E2F1, which dissociate after cisplatin. ATM/ATR inhibitor (KU-60019,5 µM) prevents cisplatin-induced decomposition of p53-p300-E2F1. P53 was co-immunoprecipitated with primary antibody and magnetic beads, whereas proteins were detected by western blot. (K) p300 remains bound to promoters characterized E2F1 motifs enriched in p53 and p300 after cisplatin-induced extrusion of p53. Genomic distribution of p53-p300 at E2F1 motifs was compared on venn diagram between untreated and cisplatin-treated cells. Table shows the number of p300 peaks overlapping p53-E2F1 sites before and after p53 extrusion. (L) DNA bound p300 is considerably enriched at p53-p300-E2F1 gene promoters after 6 h cell treatment with cisplatin. Individual p300 peak high (MACS2 score file) was normalized to median p300 peak high and plotted. p53-p300-E2F1 promoters were divided into p53 extruded and de novo recruited, further subdivided into 3 time points after administration of cisplatin.

### 3.3 p53 extrusion from p53-p300-E2F2 promoters in response to cisplatin is associated with increased gene transcription in a p300-dependent fashion

Functional analysis of the genes characterized by the occurrence of p53-p300-E2F1 at their promoters in a steady state showed a contribution to crucial intracellular processes associated with cell response to genotoxic agents (Fig. 3A). These included an *inter alia* DNA damage response, DNA repair or regulation of mitotic nuclear division. This suggested that p53 extrusion functionally contributed to cell protection from genotoxic stress. While trying to check the basic transcription level and transcriptional gene response to p53 extrusion RNA-Seq, data from untreated and cisplatin treated cells was used (Fig. 3B-D). The normalized mRNA level of genes characterized by p53 erasing and recruitment did not significantly differ (Fig. 3B), but their transcription increased substantially after the cell treatment with cisplatin (Fig. 3C). Differential gene expression indicated an almost complete overlap in both direction and intensity of the observed alteration in gene transcription after cisplatin and upon p53 deficiency (Fig. 3D). A comparable number of genes were up- and downregulated under the tested conditions (Log2FC>2, adj p < 0.05). When aligned with ChIP-Seq data on the p53 extrusion from p53-p300-E2F1 promoters, the gene subset responding to cisplatin with increased (146) and declined (45) transcription was identified (Fig. 3E). For further mechanistical study we took 2 genes from a list of 146 as the examples: *DUSP12* and *FANCI* , which are functionally linked to signaling and DNA repair, respectively. Bearing in mind that p53 erasing was followed by p300 enrichment, we tested the impact of pharmacological inhibition of p300/CBP on the transcription of genes activated by cisplatin and p53 transient silencing. In both cases, ip300/CBP completely prevented the increase in gene transcription, thereby indicating the role of p300/CBP in the observed gene activation (Fig 3F). This suggested the enrichment of p300 at the p53-p300-E2F1 promoters after p53 extrusion reversed p53-mediated gene suppression. Although p300 pre-existed with p53 at the subset of gene promoters its occurrence was insufficient for a high transcription yield.

**Fig. 3.**
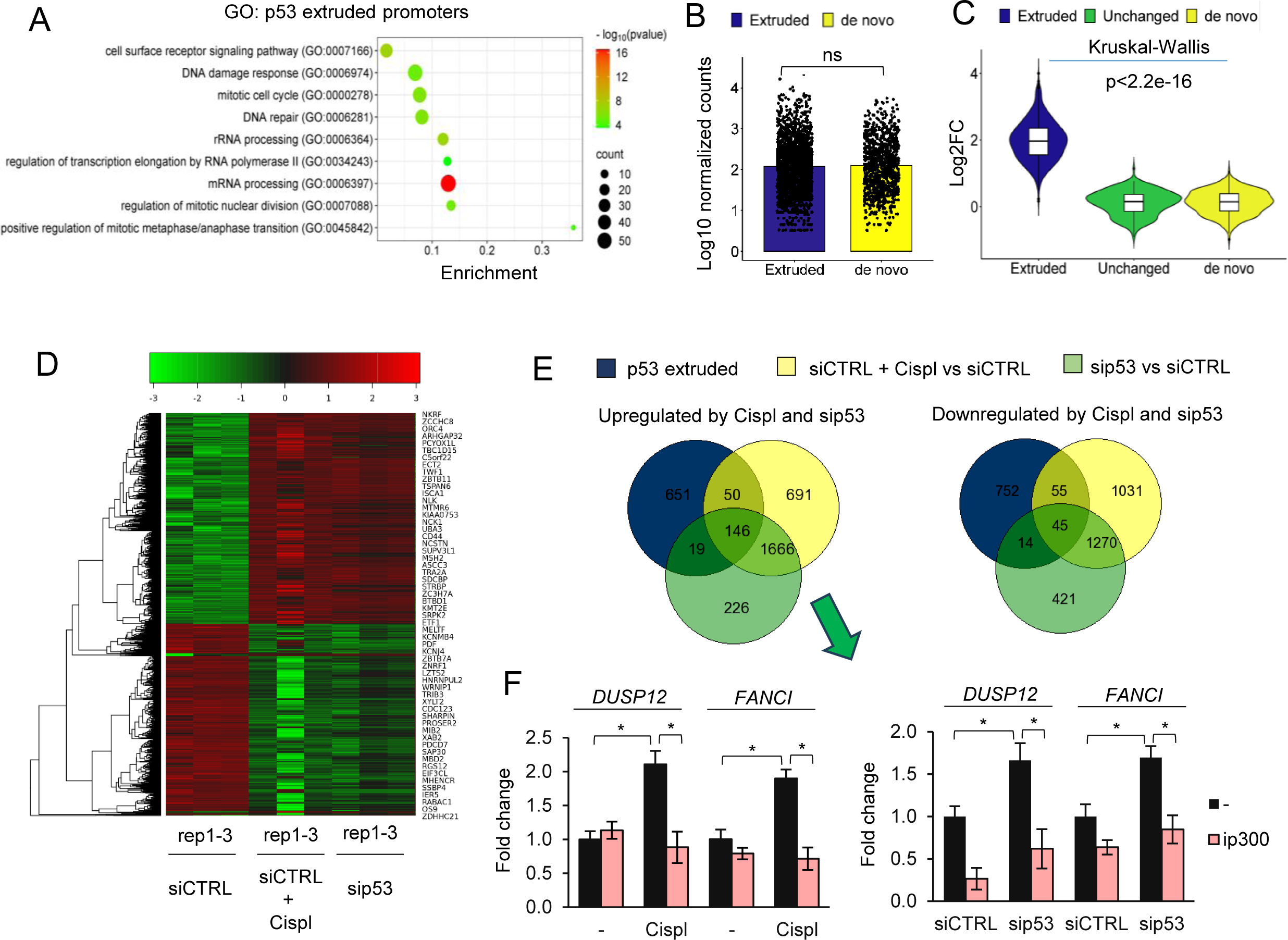
p53 extrusion from the subset of p300-bound promoters augments p300-dependent gene transcription. (A) p53 is extruded from p53-p300-E2F1 promoters, which control transcription of genes functionally linked to DNA damage response and repair, mitotic divisions and gene transcription. (B) Transcription of genes characterized by promoters extruded or de novo recruited p53 after cisplatin is comparable in untreated cells. Log10 normalized counts (counts per million) was taken for plotting and statistical analysis (t-test, ns when p>0.05). (C) Transcription of genes controlled by p300-E2F1 promoters extruded with p53 increases in response to cisplatin. Differential gene expression between cisplatin treated (24 h) and untreated cells was analyzed for promoters with p53 extruded, maintained and de novo recruited using limma-voom. Statistical analysis with Kruskal-Wallis test indicates significant differences in gene response to cisplatin among the three groups. (D) Cisplatin and transient silencing of p53 similarly affects the gene subset. Differential gene expression was carried out using limma-voom, and compared cells transfected with siCTRL, siCTRL treated with cisplatin for 24 h, cells transfected with sip53 for 48 h. Heatmap shows differentially expressed genes, when Log2FC≥1 and adj p value < 0.05. (E) 146 upregulated genes and 45 repressed genes are characterized by p53 extrusion from their p53-p300-E2F1 promoters after cisplatin. p53-p300-E2F1 promoters, which respond to cisplatin with p53 extrusion, were compared with genes up and down regulated on venn diagram. (F) p300 prevents cisplatin- and sip53-induced overexpression of DUSP12 and FANCI. P300 inhibitor (C646, 5 µM) was added for 2 h prior to cisplatin or 24 h after cell transfection with sip53, whereas gene transcription was quantified with real-time PCR.

### 3.4 P53 extruded promoters carrying p53-p300-E2F1 assembly are characterized by a higher frequency of H3K4me2 occurrence with simultaneously low transcription promoting other histone marks

In order to identify specific features of the p53 repressed promoters, publicly available datasets were used and compared with the number of overlapped peaks of selected histone modifications and proteins with p53 repressed and activated promoters (Fig. 4A). The most striking difference was observed in the H3K4 methylation pattern, particularly in the H3K4me2 to H3K4me3 ratio. Approximately 90% of repressed promoters were hallmarked with H3K4me2 and slightly higher POL2R, which suggested the pausing of this enzyme, particularly when the marker of transcription elongation represented by phosphorylated serin 2 of POLR2A, remained relatively low and comparable between the 2 promoter subsets. In contrast, p53 activated promoters were more often featured with H3K4me3. Chromatin remodelling enzymes KDM5B and HDAC1 occurred more frequently at the repressed gene promoters. Although acetylation of H3K27 was observed in all the considered promoter types, its intensity considerably varied and was lowest in p53 promoters (Fig. 4B). A similar situation was observed for di- and three-methylated H3K4. Surprisingly, the coverage of p53 activated promoters with H3K4me2 was highest in the p53 activated promoters. The transcription promoting histone marks substantially increased at the promoters of *DUSP12* and *FANCI* after cell treatment with cisplatin (Fig. 4C-D). These results suggested that p53 repressed promoters were weakly primed for transcription, with p53 extrusion by cisplatin being associated with nucleosome enrichment in transcription promoting histone marks.

**Fig. 4.**
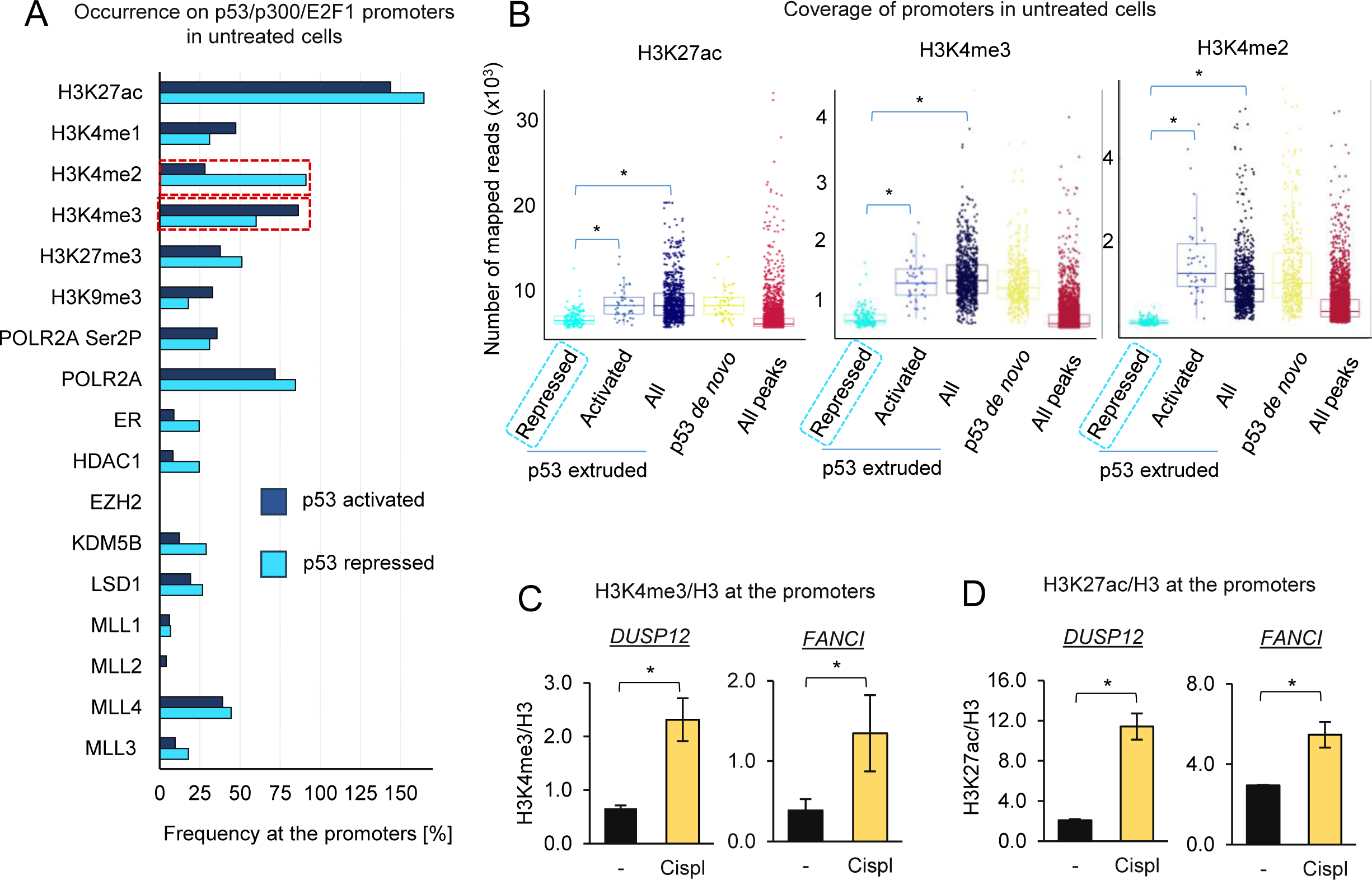
The subset of p53 repressed promoters is characterized by relatively low level of H3K27 acetylation and H3K4 trimethylation. (A)Genes activated and repressed by p53, which are controlled by p53-p300-E2F1 promoters extruded with p53 after cisplatin, are characterized by considerably distinct profile of histone marks and promoter-associated proteins in intact cells. MACS2 called peaks of considered histone marks and proteins (ChIP-Seq data downloaded from SRA) were overlapped with p53-p300-E2F1 promoters extruded with p53 after cisplatin, which were assumed as 100%. Red, cropped boxes indicate the highly variable marks. (B) Promoters of genes repressed by p53 in intact cells reveal relatively low level of transcription-promoting histone marks. The coverage of p53 peak subsets with the mapped reads of H3K27ac, H3K4me2 and H3K4me3 was calculated by bedtools MultiCovBed and compared to mean coverage of all histone mark peaks. * indicates statistical difference when p<0.05 (Kruskal-Wallis). (C-D) Cisplatin causes increased acetylation of H3K27 (C) and trimethylation of H3K4 (D) at the p53 binding sites in promoters of DUSP12 and FANCI.

### 3.5 p53 maintains KDM5B bound to p53 repressed promoters, thereby preventing p300 recruitment and conditioning low transcription yield in the steady state

Bearing in mind that the status of H3 methylation and acetylation changes after p53 extrusion, and that chromatin remodelling enzymes such as HDAC1 and KDM5B occur more frequently at the p53 repressed rather than for activated promoters, the impact of these pharmacological inhibitors on transcription of *DUSP12* and *FANCI* were compared (Fig. 5A-B). The inhibitor of LSD1 was also considered, since this enzyme erases methyl groups from mono- and dimethylated H3K4. Among the tested inhibitors, only iKDM5B phenocopied cisplatin and p53 silencing in upregulation of *DUSP12* and *FANCI* transcription with its combination with cisplatin did not further augment gene transcription. In intact cells where KDM5B overlapped p53 peaks at the promoters of *DUSP12* and *FANCI* (Fig. 5C), cisplatin caused substantial extrusion of KDM5B (Fig. 5D). To confirm the molecular link between the cisplatin-induced loss of p53 and KDM5B, enrichment of p300 and gene transcription, it was firstly confirmed that p53 deficiency at the promoters of *DUSP12* and *FANCI* caused a massive decline in the KDM5B association with chromatin, similar to cisplatin (Fig. 5E). Co-immunoprecipitation of p300 confirmed the existence of p53-p300-KDM5B complexes in a steady state, decomposing after cisplatin. Importantly, cisplatin caused a visible decline in H3K4me1 and H3K4me2 with simultaneous enrichment in H3K4me3 in p300 immunoprecipitates (Fig. 5F). KDM5B inhibition phenocopied cisplatin by increasing the level of H3K4me3 and H3K27ac (Fig. 5G-H), thereby suggesting that demethylase acts upstream of p300. This agreed with data described by Hinohara K. *et al.*[15], where the knockout of KDM5B was followed by the enrichment of H3K4me3 at p53 repressed promoters (Suppl. Figure 1J). Inhibition of p300 reverted iKDM5B-induced upregulation of the gene transcription (Fig. 5I) and provided a functional link between these 2 enzymes. Furthermore, pharmacological inhibition of KDM5B significantly increased the level of p300 at the gene promoters (Fig. 5J).

**Fig. 5.**
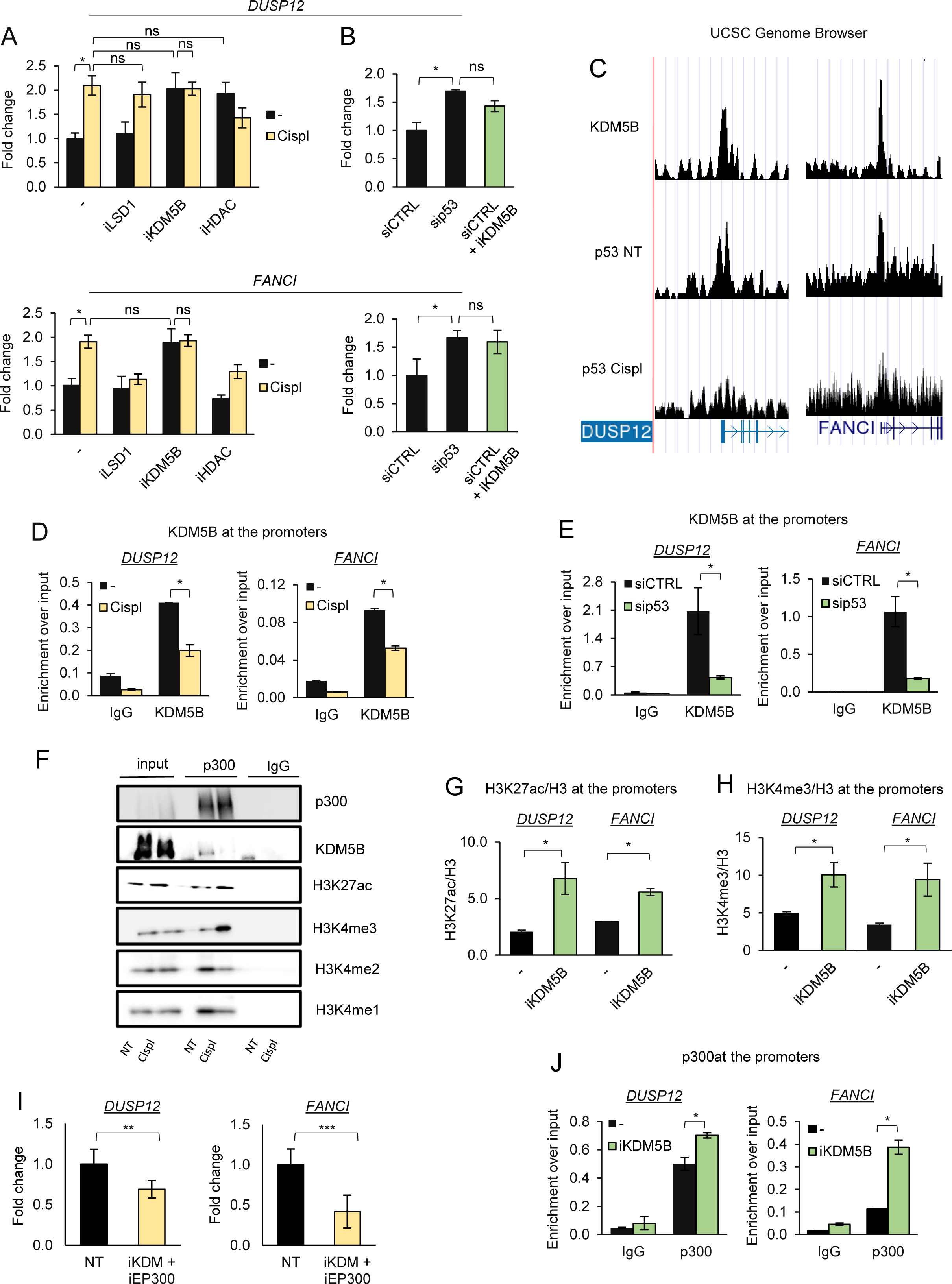
Cisplatin induces extrusion of KDM5B from the subset of p53 repressed promoters, which allows for p300-mediated overexpression of DUSP12 and FANCI. (A-B) Inhibition of KDM5B phenocopies increased transcription of DUSP12 and FANCI caused by cisplatin (A) and sip53 (B). iLSD1 (GSK-LSD1, 0,1µM), iKDM5B (PBIT, 5 µM), iHDAC (Sodium butyrate, 100 µM) were added for 2 h prior to cisplatin and 24 h after cell transfection with siCTRL. Gene transcription was measured with real-time PCR. (C) The occurrence of KDM5B and p53 (as bigwig) was visualized in steady state at the promoters of DUSP12 and FANCI. P53 track after cisplatin was aligned. (D) Cisplatin indices extrusion of KDM5B from p53 binding sites at the promoters of DUSP12 and FANCI. KDM5B association with selected genomic regions was quantified by ChIP-qPCR. * marks significant changes when p<0.05 (t-test). (E) p53 knockdown induces KDM5B extrusion from chromatin. KDM5B level at the p53 binding sites in promoters of DUSP12 and FANCI was compared by ChIP-qPCR between cells transfected with siCTRL and sip53 for 48 h. (F) Cisplatin causes decomposition of immunoprecipitable p300-KDM5B complex, but association of p300 with chromatin regions characterized by trimethylation of H3K4. Immunoprecipitation of p300 was followed by western blot detection of KDM5B, H3K4me1, H3K4me2 and H3K4me3. (G-H) Inhibition of KDM5B substantially increases the level of transcription-promoting histone marks at the promoters of DUSP12 and FANCI. Acetylation of H3K27 (G) and trimethylation of H3K4 (H) were compared by ChIP-qPCR in cells treated and not with iKDM5B (PBIT, 5 µM) for 6 h. (I) Inhibition of p300 prevents an increase in transcription of DUSP12 and FANCI caused by KDM5B. Combination of p300 and KDM5B inhibitors (C646, 5 µM; PBIT, 5 µM, respectively) was added to cells for 24 h and gene transcription was quantified by real-time PCR. (J) Inhibition of KDM5B allows for recruitment of p300 to the promoters of DUSP12 and FANCI. The level of chromatin-bound p300 was compared between cells untreated and treated with iKDM5B (PBIT 5 µM; 6h incubation) by ChIP-qPCR.

To summarise, cisplatin-induced p53 extrusion from the subset of gene promoters was associated with a decline in KDM5B occurrence allowing for recruitment of p300, the gaining of the transcription permissive chromatin structure and elevated gene transcription as shown by the examples of *DUSP12* and *FANCI*.

### 3.6 The molecular and functional interdependence between p53 and KDM5B is observed in distinct cancer cell types

To test applicability of our observation on p53-KDM5B-p300 interaction we made use of publically available ChIP-Seq datasets for human ovarian adenocarcinoma (A2780), non-small cell lung cancer (A549) and myelogenous leukemia (K562) deposited in NCBI, as well as xenograft model of A549 cells resistant to cisplatin. In contrast to MCF7 cells, the treatment of the 3 non-resistant cell phenotypes with drugs resulted in prevailing p53 recruitment to the gene promoters with relatively minor p53 extrusion from the chromatin (Fig. 6A). However, the profile of H3 marks such as H3K27ac, H3K4me2 and H3K4me3 at the p53 extruded promoters in A549 and, particularly, in A2780 resembled the promoter features observed in MCF7 cells (Fig. 6B). Also the percentage of modification-enriched promoters extruded with p53 was relatively high in A549 cells, the extent of promoter H3K4me2 was substantially higher in this promoter subset when compared to the average coverage of all p53 peaks with this histone mark in steady state (Fig. 6C). This indicates that H3K4me2 may act as a marker of p53 enriched promoters primed to p53 release upon cytotoxic or genotoxic stress. In K562 we found the subset of gene promoters characterized by the simultaneous occurrence of p53, p300 and KDM5B in proliferating cells, including the promoters of *PARP1* (Fig. 6D-E). p53, p300 and H3K4me2 peaks were perfectly centred over KDM5B enriched genomic regions (Fig. 6F-G). Since PARP1 promoter was characterized by p53 extrusion after daunorubicine, we compared the mRNA level of *PARP1* in untreated and daunorubicine treated cells. As expected, the anthracycline caused increase in *PARP1* transcription (Fig. 6H). The functional impact of KDM5B and p300 on *PARP1* transcription in steady state was confirmed by elevated mRNA level of the gene after KDM5B inhibition, and reduced after inhibition of p300 (Fig. I). The physical interaction between the 3 considered proteins was observed in untreated cells and was considerably reduced by daunorubicine (Fig. J). The similar pattern was also found in A549 cells resistant to cisplatin after their treatment with single dose of platinum drug in 2D cell culture (Fig. K). To further validate our finding, we compared p53 and KDM5B co-occurrence in the cell nuclei of tumors, which were grown in athymic nude mice and injected 6 times with PBS or cisplatin (Fig. 6L-N). Visualization of p53 and KDM5B by immunostaining and confocal imaging indicated strong colocalization of both proteins in nuclei of tumors in PBS treated mice, which was substantially declined in cisplatin-treated animals. The reduced colocalization of p53 and KDM5B was associated with their extrusion from chromatin as indicated by the decrease in their fluorescence intensity in the nuclei (DAPI positive) area.

**Fig. 6.**
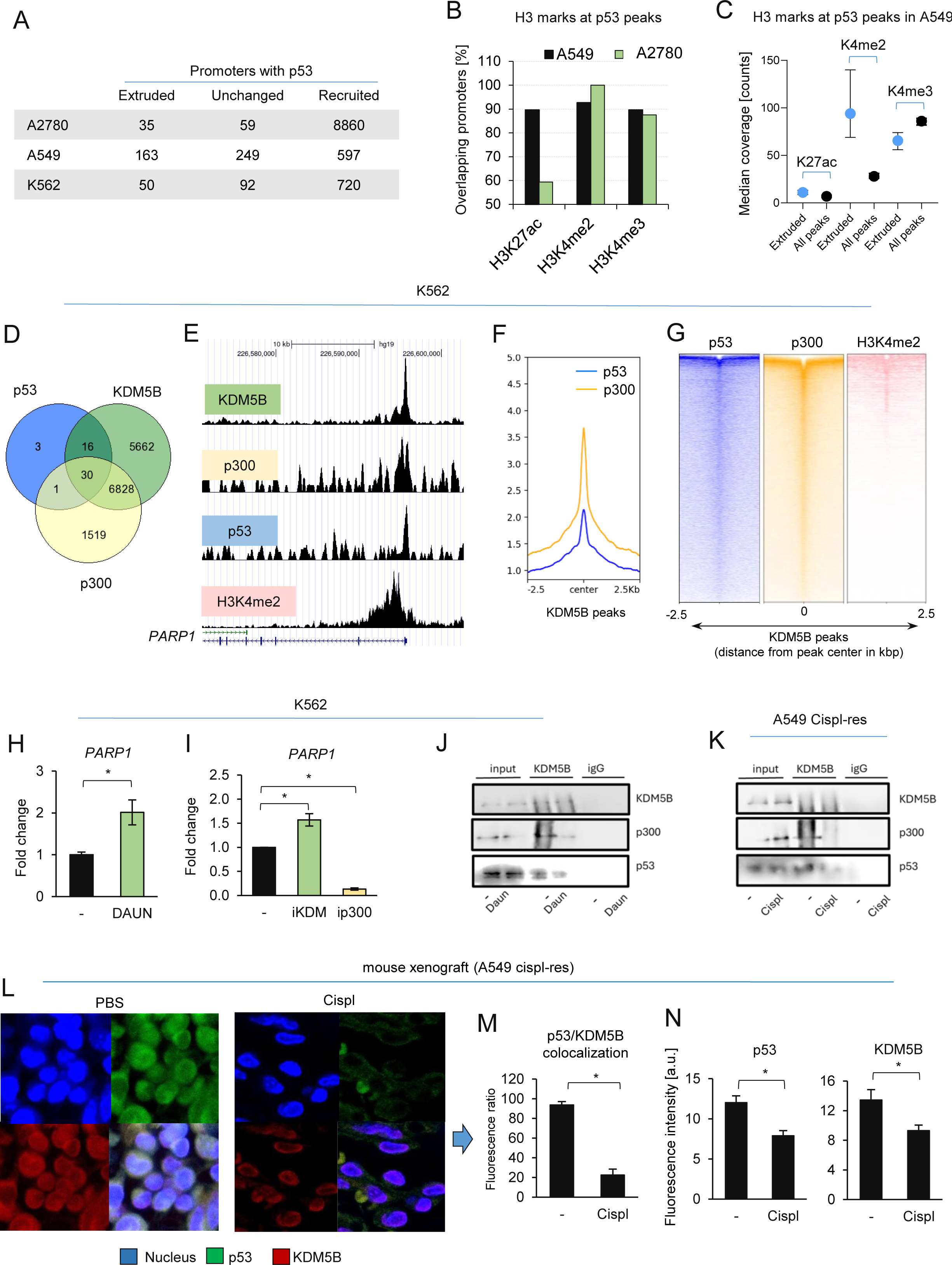
Decomposition of p53-KDM5B-p300 regulatory unit is induced by anticancer drugs in various cancer cell types. (A)The number of p53-enriched promoters in the 3 cell lines was counted by overlapping intervals of p53 peaks called by MACS2 and TSS ± 2 kbp in untreated and drug-treated cells. A270 were exposed to cisplatin, A549 to doxorubicin and K562 to daunorubicin. (B) The same approach was applied to counting promoters characterized by significant level of H3K27ac, H3K4me2 and H3K4me3 in the untreated A549 and A270. (C) The extent of promoter enrichment with histone marks was quantified by bedtools MultiCovBed on the gene promoters extruded with p53 after doxorubicin. (D) Venn diagram was used to identify gene promoters enriched in p53, p300 and KDM5B in steady state of K562. (E) Peaks of the three proteins and H3K4me2 were visualized at the *PARP1* promoter in UCSC Genome Browser. (F-G) The distribution of p53, p300 and H3K4me2 at the KDM5B peaks was analysed by plotProfile (F) and plotHeatmap (G) on the computed matrix, where KDM5B served as genomic regions in bed and mapped reads of p53, p300 and H3K4me2 in BAM served for scoring. (H-I) The changes in PARP1 transcription efficacy in K562 cells treated with daunorubicin (DAUN; 0,1 µM), iEP300 (C646, 1 µM), or iKDM5B (PBIT, 1 µM) for 24 h was measured by real-time PCR. Bars show mean ± SD and the significant difference is marked as * when p< 0.05 (ANOVA). (J-K) Immunoprecipitation of KDM5B followed by detection of p53 and p300 by western blot in K562 (J) and A549 cispl-resistant (K) exposed to daunorubicin (0,1 µM) and cisplatin (15 µM), respectively, for 6 h. (K) Tumor slices immunostained with anti-p53 (green, Alexa Fluor® 488) and anti-KDM5B (red, Alexa Fluor® 546 Conjugate) were compared between PBS and cisplatin-treated mice. (M-N) Colocalization of green and red fluorescence as well as their intensity were quantified in arbitrary units (a.u.) with Leica Application Suite X (LAS X, Leica Microsystems, Germany).

Knowing that KDM5B and p53 can co-operate in regulating transcription of some gene subsets in different cell types, we investigated whether these proteins could serve as prognostic markers of treatment resistance in cells derived from solid tumours and leukaemia’s. Transcriptome analysis of 581 cell lines treated with doxorubicin showed that low expression of *TP53* and *KDM5B* correlates with phenotype insensitive to doxorubicin (Fig. 7A). According to the ROC analysis of doxorubicin-treated cells the co-expression of these genes can be considered as a doxorubicin response biomarker (Fig.7B). The mRNA level of *TP53* and *KDM5B* correlated significantly in samples deposited in TGCA Pan-Cancer Atlas (Fig. 7C), thereby suggesting their possible intertwined role in cancer. In ER+ breast cancer low *KDM5B* expression is associated with worse patient overall survival as illustrated with kmplot.org (Fig.7D). However, no correlation between gene expression and survival was observed in patients with ovarian cancer (Fig 7G). Transcriptome analysis of ER+ breast cancer patients treated with anthracyclines (Fig. 7E) and ovarian cancer patients treated with platinum compounds (Fig 7H) indicate low *KDM5B* expression in patients resistant to therapy, whereas ROC analysis suggests that mRNA level of *KDM5B* can be considered as a anthracyclines and platinum response biomarker in patients (Fig. 7F, 7I).

**Fig. 7.**
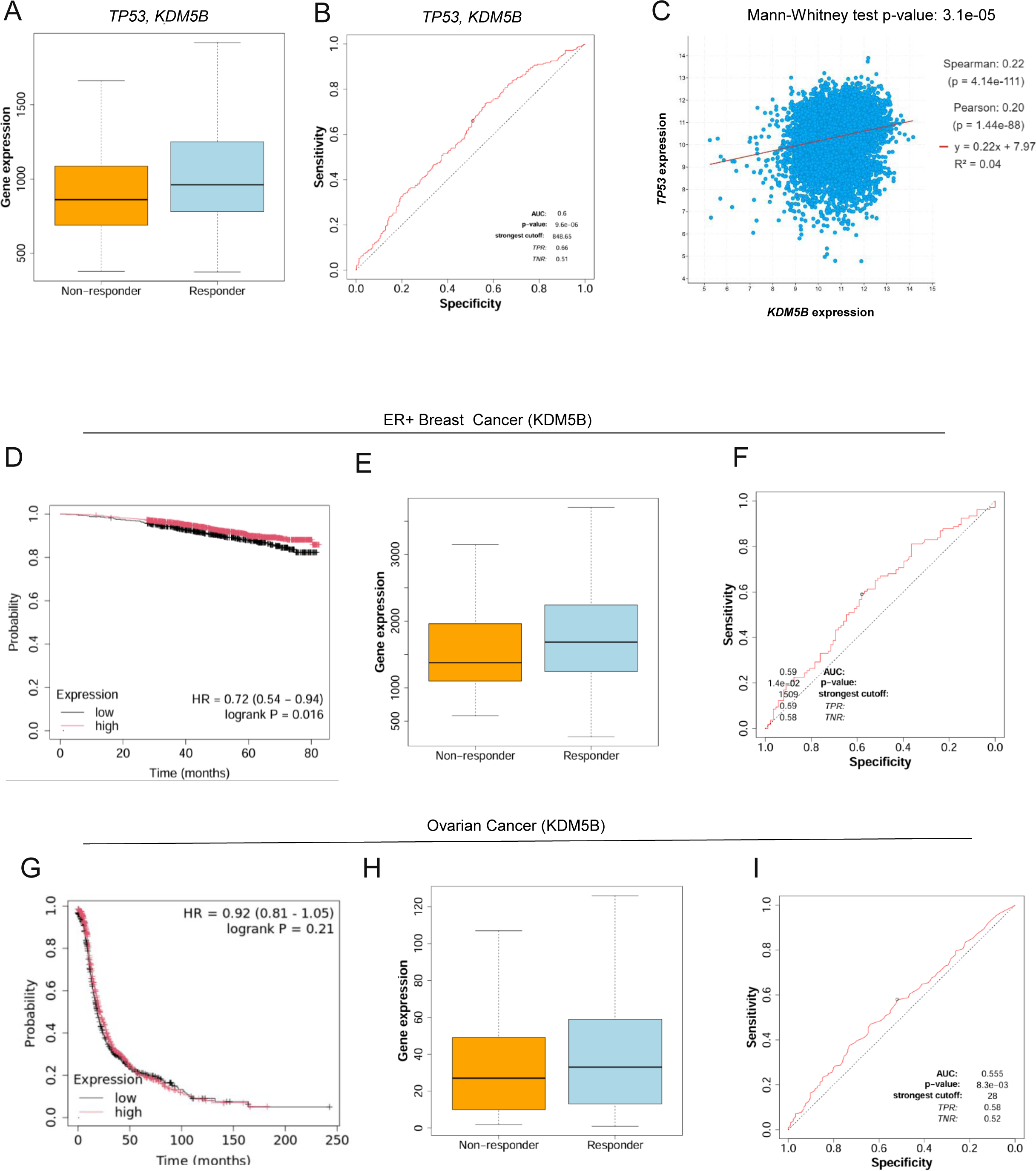
mRNA level of *TP53* and *KDM5B* can serve as prognostic markers for cancer response to drugs. (A) The dependence of solid cancers and leukemia’s cell lines response to doxorubicin therapy on TP53 and KDM5B transcription was estimated using ROC Plotter. (B) ROC curve describing the response to chemotherapy with doxorubicin in the cells characterized by simultaneously low transcription of TP53 and KDM5B C) Correlation plot with TP53 and KDM5B transcription. The underlying data were derived from all TGCA Pan-Cancer Atlas samples using cBioPortal. (D,G) Overall survival of ER+ breast cancer (D) and ovarian cancer (G) patients, which varied in KDM5B expression generated in kmplot.com. (E,H) The dependence of ER+ breast cancer patients’ response to antracyclines on KDM5B transcription (E)and ovarian cancer patients’ response to platin compounds (H) were estimated using ROC Plotter. (F,I) ROC curve describing the response to chemotherapy of ER+ breast cancer patients with anthracyclines (F) and ovarian cancer patients with platin drugs (I) in the groups of patience characterized by low transcription of KDM5B.

## 4. Discussion

P300 is one of the most frequently altered histone acetyltransferases in cancers with increased expression in some tumors and has been shown to co-activate gene transcription, together with p53. This enzyme modifies histones within proximal and distal gene regulatory elements and its activity is closely linked to H3K27 acetylation at the enhancer loci, which promotes gene expression [16]. Substantial evidence points to a critical role for p300 interaction with p53, particularly during responses to DNA damage, where p300 binds to and acetylates p53 leading to full p53 transactivation [17]. p300 binding by p53 results in the acetylation of histones and facilitated gene transcription [18]. Several factors that are activated in response to DNA damage, including CHK1 [19], SOX4 [20], MAP9 [21], PI3K [22] and RSF1 [23], were shown to act as cofactors of the p53-p300-dependent gene transcription. In this paper we provided molecular and mechanistical evidence for a p300-p53 pre-existence at the subset of gene promoters in unstressed cells and that co-occurrence of p300 with p53 was insufficient for active gene transcription. It was also reported that recruitment of p300 did not always correlate with gene activation and was occasionally associated with repression. A large number of chromatin regions enriched in p300 were not featured by canonical H3K27ac modification, indicating that histone acetyltransferase (HAT) activity at such sites was blocked, however any substantial functional explanation for this is missing [24]. In our model of proliferating cells, p300-mediated transcription activation was opposed by KDM5B, which co-occured with p53 and p300. However, the collected results did not show that the sole KDM5B extrusion from these promoters triggered catalytic activity of p300, and that under such conditions the pre-existing p300 was sufficient to enhance gene transcription. This is particularly the case when cisplatin-induced KDM5B and p53 erasure, as well as KDM5B inhibition, are followed by further p300 enrichment. Undoubtedly, the observed histone acetylation in response to the drug was catalyzed by p300.

The direct, enzymatic interaction between KDM5B and p300 has not been reported before, with these 2 epigenetic proteins being described in a negative feedback loop in the context of gene transcription rather than in regulating the activity of each other. KDM5B specifically demethylates lysine 4 of histone 3 (H3K4), often targeting trimethylated K4, which marks transcriptionally active promoters. The deregulated expression of this enzyme has been implicated in numerous cancer types, including breast, lung, liver, prostate and skin cancer [25,26]. Therefore, a direct, inhibitory impact of KDM5B on p300 activity seems unlikely, since lysine methylation of this acetyltransferase has not yet been described. The only reported arginine methylation by CARM1 in the N-terminal region of p300 inhibited the binding of the transcription factor CREB and in the C-terminal region of p300 inhibited the bimolecular interactions between p300 and the p160. Removal of this mark by PAD4 enhanced bimolecular association [27] and acted in opposition to the hypothesized p300 demethylation by KDM5B, erasing lysine methylation. However, it has been shown that lysine demethylases also have arginine demethylase activity, from studies both on histone fragment peptides, histone H4 from calf thymus (with KDM4E) and non-histone substrates (G3BP1 with KDM5C and KDM5D). Catalytic domains of all the identified human KDM5 members (A–D) have dual lysine and arginine demethylase activities in their isolated forms [28]. This suggests that further studies on p300 methylation sites and their impact on p300 activity are needed, particularly as methyltransferases are an important part of the complexes that regulate p300-dependent expression. The analyses revealed significant doxorubicin-induced enrichments of p53, H3K18ac, H3K27ac, and H3K4me1 at the enhancers, as well as Pol II, the SET1C and H3K4me3 at the gene promoters. However, p300 and SET1C seemed to act synergistically on gene transcription by the deposition of transcription promoting histone marks rather than by SET1C depositing methyl groups on p300 [29].

During DNA damage response, the ATM-mediated phosphorylation of S106 in p300 allowed interaction of p300 with NBS1, thereby enhancing DNA damage repair [30]. Much evidence has been provided on p300 phosphorylation under certain specific conditions, which opposes the impact on p300-dependent gene transcription [31–33] and stability [34,35]. In our study, activation of ATM/ATR-Chk1/Chk2-p53 did not affect p300 abundance to much degree inside the cell, but it did enhance p300 interaction with chromatin, particularly with the gene promoters characterized by motifs for E2F transcription factors. Interestingly, this was followed by a massive p300 extrusion from chromatin at a later time point after cisplatin, which suggested chromatin compaction and gene repression after the initial chromatin relaxation and gene activation. In the subset of E2F-driven promoters the existence of SWI/SNF-p300-HDAC1 complexes have been documented where the balance between p300 and HDAC1 activity was tightly controlled by cell cycle progression [36,37]. In this study one of our exemplary genes, *DUSP12,* was activated comparably by cisplatin, iKDM5B and iHDAC. This suggested that some HDAC family members may directly or indirectly co-repress some of the p53-p300-KDM5B controlled genes. KDM5B has been shown to physically recruit class I and class IIa of HDACs [38]. Approximately 50% of ∼140,000 KDM5B enriched regions overlap with the HDAC1 peaks. The cooperative action of the KDM5B and HDAC1 linked H3K4 demethylation and lysine deacetylation to provide a powerful mechanism for a rapid shutoff of actively transcribed genes [39]. Some of the HDAC family members also deacetylated p300, thereby causing its enzymatic inactivation [40]. If the gene was simultaneously repressed by HDAC, KDM5B, p53, which co-occurred with p300, the whole picture became more complex and the writing/erasing/reading processes comprised of more possible substrates.

In this study, p53 acted more as co-repressor together with KDM5B, since it supported the KDM5B binding to chromatin, but the basics of p53-KDM5B-p300 interaction remains unknown. The p53 and KDM5B did not coincide randomly, but formed an immunoprecipitable complex at a steady state with p53 occurrence being causative for KDM5B maintenance on chromatin. Previous data has shown that KDM5B represses E2F-target genes during senescence. Increased binding of KDM5B to E2F-target genes during senescence was correlated with a strong reduction of the H3K4me3 of these E2F-target genes. The suppressed transcription in senescent cells was restored and was comparable between Rb1, KDM5B and p53-knockdowns [41]. However, in that model the functional interaction between the listed proteins was observed in cell cycle arrested cells. This varied from our experimental settings, where we intentionally applied a low concentration of cisplatin to avoid the Rb-E2F-co-repressor complex formation in response to mitotic arrests. In contrast to previous reports, which focused on p53 recruitment to chromatin in response to genotoxic stress, we also showed a considerable p53 extrusion from the genome. This supported some former observations and hypotheses on the role of p53 in maintaining the transcription repressive chromatin structure [1,4], further evidenced by molecular and functional analysis. Even though p53 was thought to bind to the *CDKN1A* promoter and upstream regulatory regions, thereby activating its transcription and inhibition of mitotic divisions by the *CDKN1A* product – p21 protein, we found p53 enriched in 3 regions upstream of the *CDKN1A* transcription start site. Cisplatin caused a redistribution of p53 from location proximal to more upstream of TSS, which was associated with the decline in *CDKN1A* transcription. Of note, KDM5B has been documented as being the promoter of cancer cell proliferation by acting as a corepressor of *CDKN1A* [42]. Therefore, the dose of the drug aimed to cause DNA damage might be crucial in the transcription response of the p53-p300-KDM5B promoters characterized by E2F motifs. While low doses likely activated transcription by extrusion of KDM5B and p53, the higher doses likely repressed these genes by recruiting retinoblastoma-based chromatin remodelling complexes. However, this hypothesis requires further experimental verification.

The dose-dependent role of p53 in regulating gene transcription is particularly important when considering the functional association of p53-regulated genes. Although only a small subset of p53-KDM5B repressed promoters was focused on, the 866 promoters which responded to cisplatin with p53 extrusion were linked to intracellular processes that are crucial for *inter alia* DNA damage response and DNA repair. This protein has already been considered as a double-edged sword depending on the severity of damage. Early in the DNA damage response, p53 relays a wide range of pro-survival signals including cell cycle arrest, allowing the cells to repair the damage at various, also non-transcriptional levels. But if damage continues to accumulate, p53 promotes apoptosis or senescence. It contributes to several repair mechanisms including nucleotide excision repair, base excision repair, mismatch repair, homologous recombination repair and non-homologous end joining [43]. p53 also interacts with repair proteins like BRCA1 and DNA polymerase β, thereby enhancing the repair efficiency. As a transcription co-factor, p53 promotes the transcription of *DDB2, TP53I3*, *XRCC5* and *XPC*, facilitating DNA repair [44,45]. Previous reports also showed the p53-mediated repression of *RECQ4* during DNA damage resulting from the modulation of the promoter occupancy of transcription activators and repressors, specifically from recruitment of HDAC1 and the loss of SP1 and p53 binding to the promoter [46]. p53 also inhibits DNA repair by transactivating the base excision DNA repair inhibitory phosphatase PPM1D and by negatively regulating expression of DNA repair proteins *BRCA2*, *RAD51*, *MSH2* and *XRCC4* after genotoxic stress induced by anticancer drugs [47–49]. The activation and repression of the gene subset functionally linked to the DNA damage response by p53, allows the cell to have quick and precise activation of the most desired response pathway, that may differ in normal and cancer cells as well as strongly depend on the stimuli intensity.

## 5. Conclusions

p53 may pre-exist on the chromatin in proliferating, undamaged cells together with p300 and KDM5B and repress transcription of some E2F-driven genes. Mild genotoxic stress, which activates the ATM/ATR-Chk1/Chk2-p53 cascade, triggers extrusion of KDM5B in a p53-dependent fashion for p300 enrichment and activation of the gene transcription. P53 interaction with chromatin in a steady state may determine the cell response to damaging agents.

## List of abbreviations

CDK: cyclin dependent kinase
DSBs: double-strand breaks
HDAC: histone deacetylase
HPV: human papilloma virus
KDM: lysine demethylase
RE: response element

## Author contributions

Karolina Gronkowska: Investigation, Validation, Formal analysis, Writing – original draft, Writing – review & editing; Kinga Kołacz-Milewska: Investigation; Sylwia Michlewska: Visualization; Agnieszka Robaszkiewicz: Conceptualization, Validation, Investigation, Data curation, Resources, Funding acquisition, Project administration, Writing – original draft, Writing – review & editing.

## Data availability

Data underlying the results presented in this manuscript are provided within the Supplementary Data. Original raw data files containing sequence reads and quality scores were submitted to the NCBI Sequence Read Archive (SRA) database under the BioProject accession numbers: PRJNA1309994, PRJNA1309307, PRJNA1309881.

